# Fluidic Circuit Board with Modular Sensor and Valves Enables Stand-Alone, Tubeless Microfluidic Flow Control in Organs-on-Chips

**DOI:** 10.1101/2021.11.24.469685

**Authors:** Aisen Vivas, Albert van den Berg, Robert Passier, Mathieu Odijk, Andries D. van der Meer

## Abstract

Organs-on-chips are a unique class of microfluidic *in vitro* cell culture models, in which the *in vivo* tissue microenvironment is mimicked. Unfortunately, its widespread use is hampered by their operation complexity and incompatibility with end-user research settings. To address these issues, many commercial and non-commercial platforms have been developed for semi-automated culture of organs-on-chips. However, these organ-on-chip culture platforms each represent a closed ecosystem, with very little opportunity to interchange and integrate components from different platforms or to develop new ones. The Translational Organ-on-Chip Platform (TOP) is a multi-institutional effort to develop an open platform for automated organ-on-chip culture and integration of components from various developers. Central to TOP is the fluidic circuit board (FCB), a microfluidic plate with the form factor of a typical well plate. The FCB enables microfluidic control of multiple components like sensors or organ-on-chip devices through an interface based on openly available standards. Here, we report an FCB to integrate commercial and in-house developed components forming a stand-alone flow control system for organs-on-chips. The control system is able to achieve constant and pulsatile flow recirculation through a connected organ-on-chip device. We demonstrate that this system is able to automatically perfuse a heart-on-chip device containing co-cultures of cardiac tissues derived from human pluripotent stem cell-derived cardiomyocytes and monolayers of endothelial cells for five days. Altogether, we conclude that open technology platforms allow the integration of components from different sources to form functional and fit-for-purpose organ-on-chip systems. We anticipate that open platforms will play a central role in catalysing and maturing further technological development of organ-on-chip culture systems.

## Introduction

Organs-on-chips are microfluidic devices with integrated cultured cells that allow dynamic control over the culture microenvironment in two and three-dimensional configurations^1^. These devices allow biologists to perform assays of tissue functionality that are impossible to perform in common cell culture hardware such as well plates.^2^ There is a wide range of applications of organs-on-chips in fields like pharmacology, toxicology, stem cell biology and biomedical science^3^.

Organs-on-chips almost always include an actively perfused vascular compartment because of the essential role of blood vessels in human physiology and pharmacology^4^. The vasculature regulates transport of oxygen, nutrients and waste into and out of tissues, it is involved in the regulation of immune responses and is a major determinant of the pharmacokinetics of drugs and drug candidates^5^. Moreover, the vasculature links together all organs in the human body and is therefore essential to include in organs-on-chips when designing linked multi-organ-on-chip, or ‘body-on-chip’ systems^6^. In addition to the dynamic vascular compartments, other culture compartments of organs-on-chips are often also actively perfused, such as microbial perfusion of a gut-on-a-chip^7^, air flow in a lung-on-chip^8^ or pre-urine flow in a kidney-on-chip^9^.

Despite the numerous advantages that organs-on-chips bring in terms of dynamic control over the cell culture microenvironment using microfluidic systems, their application and operability still pose a steep learning curve for biologists^10,11^. This challenge mostly stems from the complex microfluidic set-ups that are needed to drive flow through organs-on-chips. Commercial solutions entail the use of bulky systems that are laborious to set up and that require time and expertise to implement. Additionally, the many fluidic connections and the time-consuming process of setting up these systems further increase the entry-level user barrier. Due to the numerous steps required to set up the systems failures can still occur. Typical failures include fluid leakage, infections, bubble formation, and empty fluidic reservoirs leading to dry microfluidic channels that house the cells, resulting in experimental failure. The combination of these factors are significantly limiting the implementation of organs-on-chips in end-user settings ^12^.

Multiple attempts have been made to easily integrate microfluidic cell culture devices with each other and external pumps by using either tubing or specific connectors ^13–15^. However, integrating off-the-shelve components would require additional connections and adaptors in order to interface them with the connection methods so far proposed. This hampers the adoption and integration of already existing technologies into microfluidic setups. Adopting the concept of an open technology platform for organs-on-chips, which offers developers an openly available set of interfacing standards, would enable seamless integration of commercially available components along with in-house developed devices. One such open technology platform that is being developed in a multi-institutional effort is the Translational Organ-on-Chip Platform (TOP) ^16^. Central to TOP is the use of a fluidic circuit board (FCB), which is analogous to a printed circuit board for microelectronic applications ^17,18^. The FCB serves as a physical connecting station to which microfluidic building blocks (MFBBs) can be connected. Such MFBBs can be any functional unit coupled to the FCB, e.g., sensors, reservoirs, mixers and organs-on-chips. TOP is an open platform in the sense that it facilitates collaborations and reutilization of already developed technologies by allowing connection of MFBBs on an FCB via publicly available standards. These standards are adopted from broader initiatives that aim to standardize microfluidic device design and its interconnections^17,18^. The standards describe the footprint dimensions and the grid-size to be used for inlet and outlet layout of microfluidic devices. They also describe the inlet diameter, pitch and edge distance to ensure compatibility across multiple platforms. The standards are publicly available as ISO Workshop documentation ^19–21^.

Here, we present a TOP-compatible, organ-on-chip microfluidic perfusion platform using off-the-shelve, commercial components in an integrated, stand-alone solution with a small footprint and minimal fluidic and electronic connections. We demonstrate that the concept of TOP offers the flexibility to design fit-for-purpose set-ups that can integrate both commercially available and in-house developed MFBBs. We use our setup to actively re-circulate medium through a heart-on-chip (HoC) device.

## Experimental

### Microfluidic flow control system concept and operation

The system is comprised of a FCB that interconnects the different fluidic components – MFBBs – and a control box that houses the required electronics for control. The control box houses a microcontroller, two pressure pumps, two pressure sensors and a power source required to run the system (Figure S1). The FCB acts as a printed circuit board for fluids, interconnecting all the different fluidic components in a standardized manner. The MFBBs employed in the current configuration are two solenoid valves, a flow sensor, four reservoirs and a HoC – Figure 1.b (for detailed explanation of the interconnections refer to Figure 2.a). The FCB is composed of three layers, two of which are bonded together forming the pneumatic and fluidic circuits of the system. A third bottom layer acts simultaneously as a protection case to the HoC device and as a flat resting surface that provides stability to the FCB. This layer is connected to the bottom of the FCB and the HoC can be imaged without being removed from the FCB.

**Figure 1.**
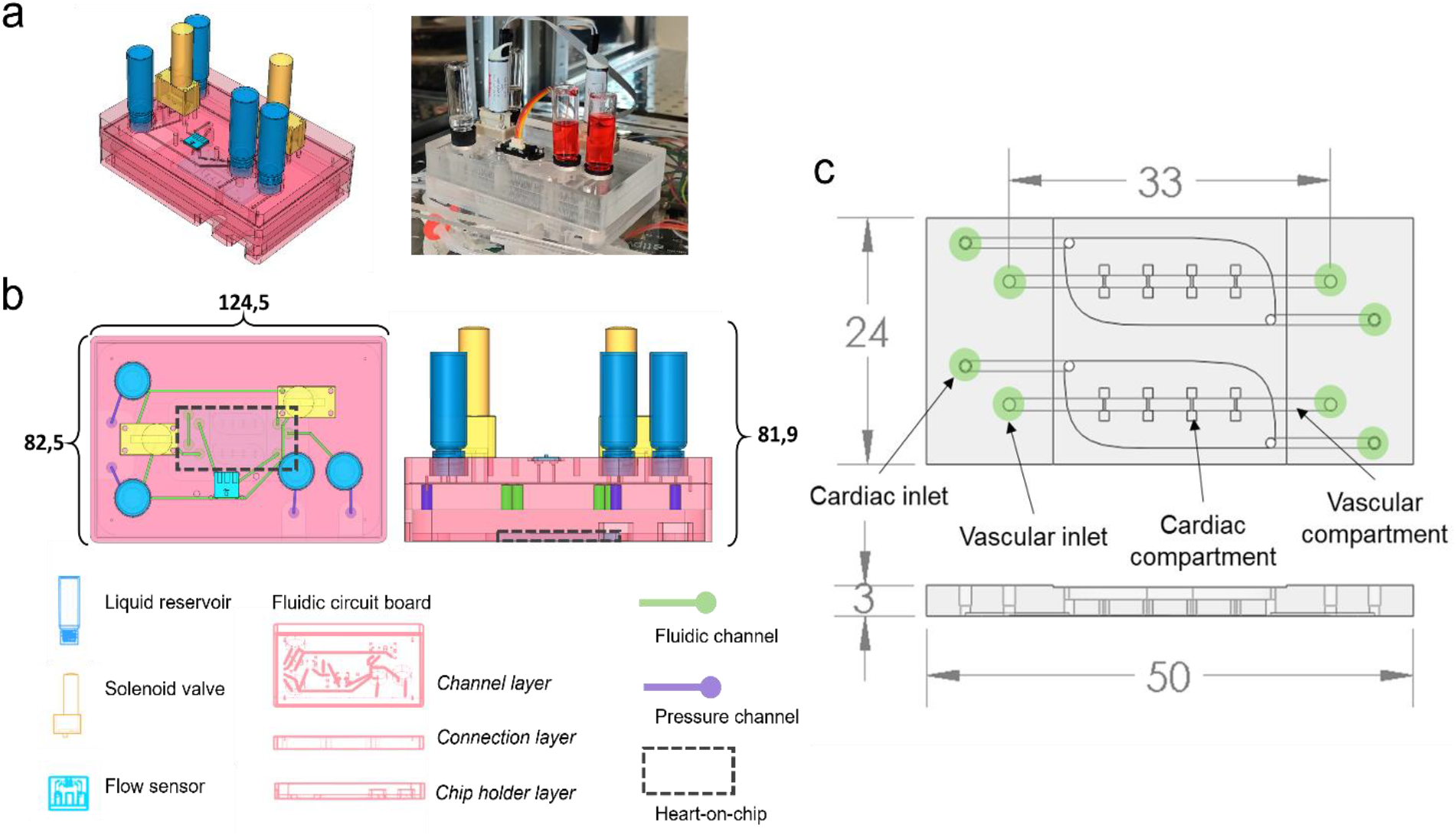
Fluidic circuit board with modular sensors and valves enables fluidic interfacing with heart-on-chip, a) Isometric view of the assembled flow control system (left, schematic; right, photograph), b) Top (left) and side (right) schematic view of the assembled flow control system with the fluidic circuit board, individual components and fluidic and pressure channels depicted. Dashed line highlights the site of interfacing with the heart-on-chip device, c) Top view (top) and side view (bottom) schematic depiction of the heart-on-chip device. The device contains two identical independent units consisting of a cardiac compartment with four cardiac chambers and a vascular compartment, separated by a porous membrane. Both the cardiac and vascular compartments have independent fluidic inlets and outlets (green). All dimensions are in mm.

**Figure 2.**
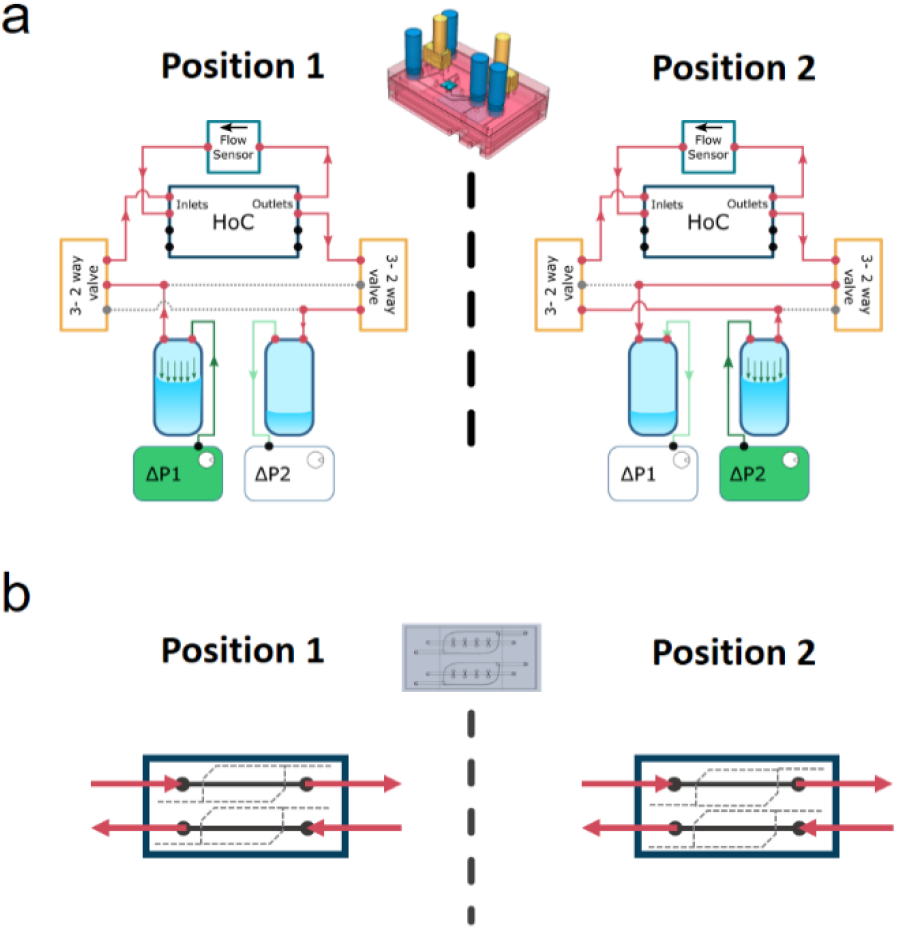
Operation of flow control system for recirculation of culture medium with unidirectional flow in the connected heart-on-chip, a) Overview of the fluidic circuit flow path between the different MFBBs and the respective flow direction in two respective Positions of the valves (left and right), b) Overview of flow direction inside the heart-on-chip device demonstrates identical flow patterns in both positions of the system.

The FCB is connected to the control box with 2 pneumatic lines that provide regulated pressurized air to the FCB and two ribbon cables that connect all electronic MFBBs on the FCB to the microcontroller in the control box. To image the HoC device coupled to the FCB with microscopy, the FCB can be decoupled from the control box and placed on a standard inverted microscope by removing the pneumatic and electronic connectors. Alternatively, live imaging with flow perfusion is also possible because of the well plate footprint of the FCB (Figure 1.b) and the portability of the control box. All the electronic components are connected to and controlled by a microcontroller programmed to regulate flow, valve position and driving pressures, as well as to monitor all these variables. For a more detailed description of the components, refer to the electronic components section below, and for the different connections among the different components to Figure S.2.

The layout of the fluidic paths in the FCB allow the unidirectional recirculation of cell culture medium by opening and closing of the valves (Figure 2). In each system position, one of the pressure controllers pressurizes a reservoir, driving the fluidic flow to one of the solenoid valves which connects to the HoC. A flow sensor is connected in series with the HoC which allows direct on-line and in-line flow measurements. One of the HoC outlets connects with another solenoid valve which leads flow to the second reservoir. The system alternates between two positions (Figure 2.a). As the system switches from one position to another, the states of the valves change, allowing unidirectional flow in the heart-on-chip channels (Figure 2.b) and reversed flow in the reservoirs.

The position and control of the volumetric flow rate is controlled by an algorithm depicted in Figure S3. The process starts by triggering a flow measurement and a comparison between the measured flow rate value against the set value. A proportional-integral-derivative (PID) controller algorithm then computes the necessary adjustment to the input pressure to drive the volumetric flow rate to the set value. The computed pressure is then sent to the active pressure controller driving the flow. The control cycle continues in a repetitive loop until the set time has been reached. This cycle time is required to avoid completely emptying the pressurized reservoir and drying the microchannels. The elapsed time of the control cycle loop is checked with an ‘IF’-condition programming statement. If the statement is true, the current pressure controller is set to 0 and the valve positions are changed. The cycle then starts over until the cycle time is reached again. If not, these instructions are skipped, and another cycle starts.

## Materials and Methods

### FCB fabrication

The FCB was composed of two casted 10 mm PMMA plates (Altuglass, France) where all connecting channels and fittings for Luer-slip connectors were milled with a CNC micro mill (Datron Neo, Datron AG). The milling models are available as downloadable files at the GitHub repository.^22^ The channels in the FCB have a rectangular cross-section of 1 mm × 0,5 mm (width×height). Inlets were placed in a 1.5 mm grid following published standards ^19–21^. The footprint dimensions of the FCB were reduced by 3.7 % to those of the ANSI standard well plate ^23^ footprint to accommodate for tubing connections when the FCB is mounted on a microscope stage. This reduction in size does not affect the FCB compatibility with common well plate mounting hardware. In terms of height, however, the FCB is currently 9.5 mm taller than the well plate standard.

After milling, both layers of the FCB were cleaned using industrial cleaning wipes (Adolf Würth GmbH & Co), rinsed under running deionized water and blown dry with compressed nitrogen. Subsequently, both slabs were rinsed with 100% ethanol and isopropyl alcohol and again blown dry with nitrogen. A solution of acetone in pure ethanol at a volume ratio of 1:10 was added on top of the connection layer slab. The complementary channel layer slab was then pressed onto the connection layer slab and aligned using 1 mm diameter and 6 mm length alignment pins (DIN 7 - ISO 2338, Thormas, ERIKS BV, Netherlands) inserted in 1 mm pre-drilled holes using the previously mentioned micro mill. The assembled FCB was then pressed at 1 kN at 55°C using a hydraulic press (model 3889, Carver Inc.) during 5 minute intervals and checked for the absence of colored interference fringes. If interference fringes were seen, more acetone solution was added through the edges of the FCB by capillary action until the interference fringes disappeared and pressed again for another cycle. The process was repeated until the FCB was bonded – typically requiring 2 to 3 cycles. The FCB was then brought to room temperature (approx. 25°C) using the water-cooling function of the press.

### Commercially available microfluidic building blocks

All the electronic components, including the commercially available MFBBs used are summarized in Table 1. The individual MFBBs are briefly described below.

**Table 1.** Summary of the number of electronic components that are included in the FCB and the control box.

| Component | Component model | Qty. |
| --- | --- | --- |
| Microcontroller - MCU | Espressif ESP-32<br>Wemos Lolin32 | 1 |
| Pressure Sensor | Honeywell ABP Series | 2 |
| Flow Sensor | Sensirion LPG10 | 1 |
| Valve controller | Adafruit breakout board - DV833 | 1 |
| Solenoid Valve | The Lee Company - LFR model | 2 |
| Pressure Pump | TTP Ventus - Series micro-pumps | 2 |
| Voltage regulator | Buck converter Mini 360 (MP2307) - Breakout board | 3 |

### Reservoirs

Glass reservoirs (Sigma-Aldrich, inc) were connected to the FCB via two blunt needles (GA 21, 0.5” and GA 23, 1.5”, Nordson EFD). The needles were tightly inserted in the FCB and rapid gel glue (Pattex, Henkel) was then applied around the base of the needles to avoid removal and any possible leakage.

### Solenoid valves

Two solenoid valves (LFRA-1252170D, The Lee Company) were connected to the FCB by face-mounting them following the manufacturer’s instructions. The valves were electrically connected to the control box using the provided connectors and a ribbon cable. More precisely, the valves were connected to a 2 H-bridge motor control breakout board (DRV8833 DC/Stepper Motor Driver Breakout Board, Adafruit) controlled by the microcontroller.

### Flow sensor

A face mounted flow sensor was employed for flow control (LPG10-1000, Sensirion GmHb). The sensor has an accuracy of 5% of the measured flow in the range from 0.05 to 1 ml/min. For flow rates below 50 μL/min, the accuracy is 2.5 μL/min. The flow sensor was mounted and operated according to the manufacturer’s specifications. Briefly, the profile of the sensor and its associated o-rings was milled in the FCB and two holes for M2.5 screws were thread milled using the previously mentioned micro mill. The sensor was then clamped in place using the company’s provided mounting piece and screwed into place based on the manufacturer’s instructions. The electrical connections were made for communication with the microcontroller based on the company’s application note. Publicly available scripts were adopted for communication between the microcontroller and the flow sensor ^24^, using a cyclic redundancy check (CRC) to avoid data corruption.

### Heart-on-chip fabrication

#### Chip design and Fabrication

The HoC device is comprised of two independent vascular compartments which are each connected to their own respective set of four cardiac compartments. The cardiac and vascular compartments are fluidically connected through a 10 μm thick polyester (PE) porous membrane with 8 μm diameter pore-size (GVS Life Sciences, USA). The cardiac compartments consist of dumbbell-shaped wells similar to what has been reported previously ^25^ in which 3D cardiac tissues were fabricated. The cardiac dumbbell-shaped wells are 3.2 mm in its longest axis, where the two squares at the edges are 1 mm2 connected by a 1.2 mm by 0.25 mm shaft and a total depth of 1.5 mm. Above the cardiac tissue compartments lies the cardiac reservoir that can hold approximately 200 μL of medium when reversibly closed by a glass seal. The vascular compartments are straight rectangular-shaped channels with a cross-section of 1.2 mm width, 0.1 mm height and a length of 33 mm. Following published standard guidelines,^19^ inlets were placed in a grid of 1.5 mm in the x axis of the HoC but not on the y axis due to the spatial constraints imposed by the employed commercial coverslips – the shortest side of the HoC.

Devices were fabricated using an injection molding-like technique using a pair of negative replicate molds similar to a technique previously reported ^26,27^. The HoC and the two respective negative molds were designed using SolidWorks (Dassault Systèmes, France). A CNC machine (Datron Neo, Datron AG) was used to micro mill the molds using 8 mm thick casted poly(methyl methacrylate) (PMMA) (Altuglass, France) as a stock material. Dimensions of the molds were verified by optical microscopy using a polydimethylsiloxane (PDMS) (Sylgard 184 Silicone elastomer kit, Dow corning, USA) casted replica. The HoC was fabricated by sandwiching 1 mm wide PE membrane strips perpendicularly aligned to the channel and between the two negative molds. The membrane strips were then fixed in place by 3 mm wide and 2 cm long double-sided tape strips (3M, USA) placed parallel to the longest side of the HoC. The two PMMA molds were brought together and clamped with two N46 neodymium nickel-plated square magnets rated with ca. 58.8 N force (Webcraft GmbH).

PDMS was mixed in a 1:10 (wt:wt) polymer base to crosslinker agent ratio, degassed and injected with a 12 ml syringe (BD plastics) into the assembled mold. Air bubbles formed during PDMS injection were allowed to escape during a 30-minute waiting period at room temperature followed by a 65°C overnight incubation in an oven for PDMS cross-linking. After curing, molds were disassembled, PDMS devices were peeled off the molds and the edges were trimmed with a scalpel. Inlets for the vascular compartments were 1 mm diameter and were integrated in the mold design.

Glass coverslips with dimensions 50×24×0.15 mm (Menzel Glazer, Thermo scientific) were spin-coated (1500 rpm, 30 s, 1000 rpm/s, Spin150, Polos, The Netherlands) with a liquid PDMS polymer/crosslinker mix and cured at 65°C overnight.

HoC and PDMS-coated coverslips were simultaneously exposed to air plasma (50 W) for 40 seconds (Cute, Femto Science, South Korea), bonded and incubated at 65°C for at least 3 hours to enhance bonding.

#### Heart-on-Chip surface chemical functionalization

Fully assembled devices were silanized to enhance cell adhesion to the vascular compartment. Devices were first exposed to air plasma (50 W) and followed by filling channels with an aqueous solution of (3-Aminopropyl)triethoxysilane (APTES) (Sigma Aldrich, Germany). Devices were then incubated for 1 minute at room temperature, submerged in 100% ethanol and channels flushed with pure ethanol. Devices were then air blown dried with nitrogen and incubated in an oven at 65°C to complete ethanol evaporation.

After incubation, UV-sterilization was performed during 30 minutes in a laminar air flow cabinet (Telstar, The Netherlands). Vascular compartments were coated with a rat tail collagen type I (VWR) solution of 0.1 mg/ml in Dulbecco’s phosphate-buffered saline (DPBS, ThermoFisher, USA), and incubated for 30 minutes at 37°C in a humidified incubator. The collagen solution was then flushed out of the channels, after which they were filled with DPBS until cell seeding.

### Heart-on-chip connection to FCB

The assembled HoC with seeded cells (below) was connected to the FCB by male Luer-barb 1/16” connectors (Cole-Palmer). Care was taken not to introduce air bubbles by employing droplet contact between the FCB and the HoC. To accomplish this, first a dummy device – *i.e*., a sterile HoC without any cells - was used to run culture medium through the channels and fill all fluidic paths of the FCB. The dummy device was then removed and drops of medium were placed right on the inlets of the HoC. The Luer-barb connectors from the FCB were then visually aligned with the HoC inlets and pushed onto it, effectively sealing the connection. The excess medium was then aspirated. All these steps were performed inside a laminar flow hood to reduce chances of contamination.

### Control box fabrication and software

The control box was fabricated from sheets of PMMA glued together, and contained two pressure controllers, one microcontroller unit, and three buck converters for voltage control. All the electronic components used in the control box are summarized in Table 1.

Communication between the flow sensor with the microcontroller was stablished using an Inter-Integrated Circuit (I2C) protocol, while the communication with the pressure controllers was performed using a universal asynchronous receiver-transmitter (UART) protocol. The solenoid valves were controlled using a 2 H-bridge breakout board controlled by the microcontroller.

The microcontroller was programmed using the PlatformIO integrated development environment (IDE). The code and schematics can be found at the GitHub repository. ^22^

Two pressure lines from the pressure controllers in the control box were connected to the FCB by PTFE tubing with ID 1.6 mm and OD 3.2 mm (Sigma-Aldrich), and connectors (Luer Luer-slip to 1/16” barb (Cole-Parker)).

### Fluidic characterization

All measurements were performed using the electronic components previously mentioned and all data were recorded using the USB serial communication of the microcontroller coupled to a Python script used for data logging and analysis.

The hydraulic resistance characterization, was run in SolidWorks Flow Simulation module (Dassault Systèmes, France), using water at 25°C, with no gravity and with no slip condition.

The frequency domain analysis was performed applying an up-chirp – a linearly increasing frequency sweep – from 0 to 1 Hz at a 0.002 Hz/sec. The generated setpoint and the measured flow rate signals were then processed using the fast Fourier transform function – rfft - from numpy^28^, divided and filtered using the Savitzky-Golay filter included in SciPy^29^ with a first order polynomial and a window size of 9 data points.

### Cell differentiation and culture

Cardiomyocytes (CMs) were derived from a human pluripotent embryonic stem cell line (hESC) with two reporter genes for the transcription factor NKX-2.5 and the sarcomeric protein α-actinin, fluorescently tagged with GFP and mRuby, respectively and differentiated as previously reported ^30^. Briefly, hESCs were seeded at a density of 25×10^3^ cell/cm^2^ on Matrigel-coated 6-well plates in Essential 8 medium (ThermoFisher, USA) on day −1. At day 0, mesodermal differentiation was initiated by addition of Wnt activator CHIR99021 (1.5 μmol/L, Axon Medchem 1386), Activin-A (20 ng/mL, Miltenyi 130–115-010) and BMP4 (20 ng/mL, R&D systems 314-BP/CF) in Bovine Serum Albumin (BSA) Polyvinylalcohol Essential Lipids (BPEL) medium. At day 3, Wnt was inactivated by adding XAV939 (5 μmol/L, R&D Systems 3748) in BPEL. ^31^ Cell cultures were refreshed with BPEL on day 7 and 10 after the start of differentiation until differentiation was completed. At day 13 CMs were then frozen in medium C containing 50% Knock out serum, 40% BPEL, 10% DMSO and 1x RevitaCell and stored in liquid nitrogen. Before CMs were thawed, 6-well plates were first coated with vitronectin (5 μg/mL, Thermo Fisher, USA) followed by a coating with 10% fetal bovine serum (ThermoFisher, USA) in DMEM (Sigma, USA) for 1 h and 30 minutes at 37°C in a humidified incubator, respectively. CMs were thawed and plated on coated 6-well plates at a cell density of 1×10^5^ cell/cm^2^ and cultured in cardiomyocyte maturation medium^32^ supplemented with 100 nM triiodothyronine hormone (T3) (Sigma-Aldrich), 1 μM dexamethasone (Tocris) and 10 nM LONG R3 IGF-1 (IGF, Sigma-Aldrich, USA) (CM+TDI) for 3 to 4 days prior to the 3D tissue fabrication.

Human umbilical vein endothelial cells (HUVEC) (pooled-donor, Lonza) were sub-cultured on 0.1 mg/ml rat tail collagen type I (rat tail collagen I, ThermoFisher) T-75 coated flasks and passaged at about 80% confluency. Cells were used up to P5.

Human adult cardiac fibroblasts (cFBs, Bio-Connect) were subcultured following manufacturer’s instructions in T-75 culture flasks with FGM-3 medium (Bio-Connect). cFBs were used between P4 and P7.

CMs, cFBs and HUVECs were washed with DPBS and dissociated with 1× TrypLE select (ThermoFisher, USA) for 3 min. in a humidified incubator at 37°C. TrypLE was diluted with DMEM supplemented with FBS for CMs, FGM-3 for cFBs and supplemented EGM-2 for HUVECs. Cells were then centrifuged at 240×*g* for 3 minutes and the supernatant was aspirated. The percentage of CMs in the resulting differentiated population was quantified by flow cytometry (MACSQuant VYB flow cytometer, Miltenyi Biotech) using the fluorescent reporter markers expressed by these cells.

### 3D cardiac tissue fabrication

Cardiac tissues were formed in the microfluidic chip by seeding CMs and cFBs inside a fibrin hydrogel in the microcompartments. After dissociation, CMs and cFBs pellets were resuspended at a concentration of 5.8×10^7^ cells/ml in CM+TDI supplemented with horse serum (CM+TDI+HS) and FGM-3, respectively. Thrombin from bovine plasma (ThermoFisher, USA) was mixed into the cell suspension at 1:300 ratio with a final gel concentration of 0.67 U/ml. Both cell types were added in an ice bath to a hydrogel mix with the following composition: fibrinogen from bovine plasma (2 mg/ml, Sigma-Aldrich); Matrigel (1:10 V/V, Corning); 2 times concentrated CM (2x CM); and aprotinin from bovine lung (1:150, Sigma). The final cell concentration was 4.0×10^7^ cells/ml and the CMs to cFBs ratio was 1:10.

Four microliter of pre-polymerized gel were pipetted onto each dumbbell shape and incubated at room temperature for 10 minutes to allow hydrogel polymerization. CM+TDI+HS was then added to the cardiac compartment, and 3D tissues were cultured at 37°C and 5% CO2 in a humidified incubator for 10 to 11 days. Medium was changed every 3 days.

HUVECs were cultured in the microfluidic chip by seeding a highly concentrated cell suspension followed by continuous medium refreshment. After dissociation, HUVECs pellets were resuspended at a concentration of 5.0×10^6^ cells/ml and seeded in the collagen-coated channels of the HoC. The device was flipped upside-down and incubated for 30 minutes in a humidified incubator at 37°C to promote cell attachment on the ceiling of the vascular compartment. After incubation, non-attached cells were removed by flushing with fresh medium. Pipette tips containing 100 μl of medium were mounted on the inlet and outlet of each endothelial channel. The seeded chips were placed on a custom-made rocking platform with a 35° tilting angle with a 30 s cycle in a humidified incubator at 37°C for 3 days until confluence was reached. After confluence was reached, chips were then connected to the FCB and perfused through the vascular compartments at 100 μL/min for 5 days.

## Results and Discussion

### Modularity, integration and reduced footprint

The fluidic control system is a stand-alone modular solution for fluidic flow control of organs-on-chips. The system is comprised of two units: the FCB and the control box. The FCB routes the fluids among the different MFBBs and combines them into an integrated modular system (Figure 1). The different MFBBs were either face mounted or connected with Luer-slip barb connectors to the FCB, greatly reducing the number of connections to be made to interface all the MFBBs as well as the amount of tubing required. In total, 8 MFBBs were connected to the FCB including 4 fluidic reservoirs, a flow sensor, 2 solenoid valves (on the top side of the FCB) and one organ-on-chip (at the bottom side of the FCB). The electronic components of the FCB, *i.e*., the flow sensor and the solenoid valves, were connected to the control box using a ribbon cable to transmit sensor data to the microcontroller and power to the solenoid valves. Up to 4 pneumatic lines can be connected to the FCB where 2 connections are required to control flow in the vascular compartment and the other 2 for the cardiac compartment. By sealing the pneumatic connections to the cardiac compartments, no flow is present in the cardiac compartments. Fresh medium can only reach the cardiac compartment through mass transport from the vascular compartment. This was the mode used in this study.

To ensure compatibility of the FCB with microscope stages with a well-plate format, the footprint was reduced by 3.3 mm in length and 3 mm in width to accommodate the pneumatic tubing coming from the sides. The size reduction does not affect the FCB fitting on the microscope stage used in this study. Still, the FCB is much taller than common well-plates and not all microscopes may accommodate the current design. Advances in valve technologies, however, are bringing to market much smaller valves that use different principles of actuation, *e.g*. by shape memory alloys.^33^ The reduced footprint of such valves could allow further reduction of the FCB height and better integration in most microscope imaging systems.

The control box houses the electropneumatic modules used to control fluidic flow in the FCB. Inside the control box were the different electronic components that were not in contact with liquids, *i.e*., the pressure controllers, power source and the microcontroller using off-the-shelf components. Although the use of this format with separate components increased the control box footprint, it allows easy replication of the system without the need of custom-made electronic hardware such as printed circuit boards.

In contrast with many other organ-on-chip flow control systems, the reduced footprint of the system enables us to place the full system (FCB and control box) inside an incubator, reducing the chances of bubble formation and pH buffer alkalinization which can occur when pumping equipment is placed outside the incubator.

### Fluidic characterization

The characterization of the flow control in our system was performed in terms of fluidic resistance and two flow profiles: constant and pulsatile. Moreover, flow volume tracking was also assessed to ensure reliability of media recirculation. Control over the dynamics of medium flow in the system (including connected organs-on-chips) is important from a physiological perspective. Blood vessels are exposed to a pulsatile blood flow pattern that gradually dissipates downstream of the heart. It is of interest to also apply such flow patterns to endothelial cells *in vitro*. Moreover, for some organs-on-chips, it may be needed to gradually increase flow rates over the course of days, in order to allow cells to first form a monolayer before being exposed to higher flows.

### FCB hydraulic resistance

To ensure passive behavior of the FCB, its channels were designed to produce negligible hydraulic resistance when compared to that of the HoC. The negligible hydraulic resistance of the FCB simplifies the integration of the different MFBBs ensuring these are the source of the major hydraulic resistance in the system. As depicted in Figure 3.a, the main hydraulic resistor of the FCB is the HoC device. By adopting this design philosophy, the electric circuit analogue representing the fluidic circuit of the system is greatly simplified, similar to how the resistance of tubing can typically be ignored in other flow control systems for organs-on-chips.

**Figure 3.**
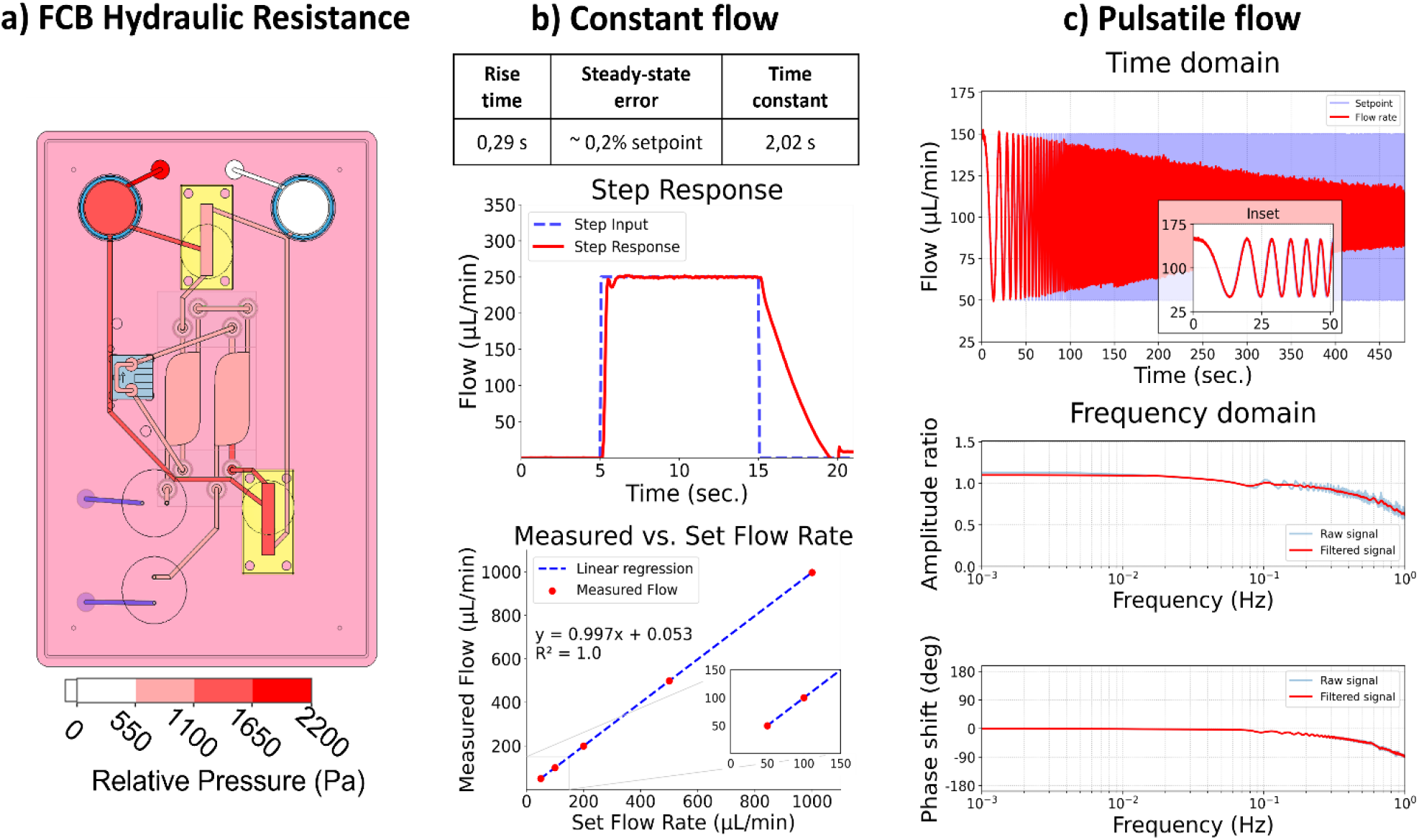
Fluidic characterization of the FCB and the implemented control system, a) Top schematic view of the FCB demonstrating the relative pressure loss throughout the fluidic path. The major pressure loss occurs at the heart-on-chip device while pressure throughout the fluidic paths remains constant, b) Characterization of the system was performed by assessing the rising time (how fast the system reaches the set value), the steady-state error (how close the flow is to the set flow) and the time constant (the time required by the system to return to zero). In the bottom graph the set flow rate is plotted against the measured flow rate along with the linear regression curve demonstrating the linear response of the system, c) Pulsatile flow characterization of the system, in the time and frequency domain. In the time domain section, a frequency sweep of the setpoint (blue) and the measured flow (red) are depicted demonstrating the stable response of the system at low frequencies – inset. In the frequency domain section, the amplitude ratio and the phase shift between the setpoint and measured flow rate are depicted.

### Constant flow

For the constant flow profile, the main challenge lies in having a unidirectional flow perfusion in the HoC channels while medium flows back and forth between the media reservoirs. A common approach is to use recirculation valves normally used in chromatography to route medium flow throughout the setup. Culture medium flow is thus stablished across two reservoirs by switching the valves’ positions and a control loop that diligently re-establishes the preset flow after the valve switch.

In order to characterize the responsiveness and control capabilities of the system, a step response analysis was performed. In this analysis, several parameters are characterized to gain insight into the system. The time span required by the system to react to an input change and go from 0.1 to 0.9 of the set input value is denominated as the ‘rise time’. The ‘time constant’ of a system is defined as the time required by the system to reach 63.2% if the measured rate would be kept constant. The time span required for stabilization of the flow is characterized by the ‘settling time’ of the system, *i.e*., the time required by the system to reach a steady state. Another important parameter is the ‘steady-state error’, which describes the difference between the set flow and the measured flow. In this system the rising time was 0.29 s, the settling time was 0.94 s and the average steady-state error was within 0.2% of the set value of 250 μL/min: 249.63 ± 0.82 μL/min (Figure 3.b). Overall, the system was able to reliably perform for the intended application.

The lowest flow rate to be used in the system is 25 μL/min, due to the flow sensor accuracy limitations. Based on the specifications of the flow sensor, lower flow rates would have an error higher than 10%. For our purpose in the current study, a minimum flow rate of 25 μL/min is acceptable, but if lower flow rates are needed, another flow sensor with a better suited range should be integrated to increase the accuracy of the system. The system was able to precisely stablish a 5 μL/min flow rate demonstrating that the limiting factor is in the accuracy of the selected sensor and not in the pressure controllers or its implementation. It is thus important that the specifications of each of the different MFBBs are evaluated prior to the design of the FCB and the integration of respective components. The maximum flow rate is also dependent on the flow sensor accuracy, and therefore attention must be paid to the specifications of the flow sensor employed.

With the present configuration of the system, it is possible to reliably apply flows between 25 and 1000 μL/min. Perfusion of the HoC with these flow rates lead to local shear rates in the vascular compartment between 208 s^-1^ and 8333 s^-1^, respectively. These shear rates cover the entire physiological and pathological range as found *in vivo*, from veins to stenotic arteries ^34^. This enables application of the system in studies of *e.g*. immune cell infiltration ^35^ and plaque formation in atherosclerosis ^36^.

### Pulsatile flow

Preliminary characterization of the system for pulsatile flow was performed within a physiological frequency range, from 0 Hz to 1 Hz. This was accomplished by using a frequency sweep sinusoidal signal as the set flow rate and analyzing the resulting measured fluid flow (Figure 3.c, time domain). Increasing the frequency is expected to result in amplitude loss, *i.e*., the peak set flow would not be reached which translates into lower shear stress values in the HoC than the ones intended. As depicted in figure 3.c in the frequency domain, the amplitude loss gradually increased as frequencies increased to 1 Hz, at which point the loss was 36%, accompanied by a negative phase shift of 91°. The seen artificial gain at low frequencies, below 0.2 Hz, in the amplitude ratio graph derives from the reduced capturing time and consequent inability to resolve low frequencies with the selected capturing time. Overall, the amplitude loss and phase shift were minimal for frequencies below 0.2 Hz, evidenced in the time domain graph inset.

Phase shift and amplitude loss stem primarily from physical effects and the implementation of the automation algorithm. Physical effects include the system pneumatic capacitance and an interplay between viscosity and transient inertial effects that affect how fast the fluid can move in the system in response to pneumatic action. In terms of the algorithm implementation, the PID tuning parameters affect how the microcontroller adjusts the pneumatics to meet the required flow, *i.e*., the control loop. Several techniques and software are available for tuning such parameters, but unfortunately these parameters must be tuned for each specific system and flow regime independently. Automation of this step is important to enable a user-friendly approach and it is subject to further development.

The system in its present form can apply pulsatile flows that mimic physiological conditions in a heart-on-chip in terms of frequency response. However, the amplitude of the pulsatile flow will be affected as depicted by the amplitude loss and should be characterized for each individual system. This result highlights the importance of integrating auto-tuning control system algorithms that facilitate the use of such platforms by non-experts. Additionally, lowering the system’s capacitance by using smaller reservoirs or reducing the amount of air inside them would aid in increasing the system’s performance at higher frequencies.

### Media recirculation

Stimulation of cultured cells with shear stress requires considerable amounts of culture medium due to the relative high flow rates applied. Moreover, the excessive medium volumes that cells are exposed to may dilute auto- and paracrine signaling molecules that cells may secrete. Recirculation of medium then becomes imperative, both for economical and biological reasons.

The flow recirculation cycle in the system can be controlled in two different ways, either by fixed intervals or by the volume pumped. These modes can also be combined for redundancy and to avoid emptying either of the reservoirs. In the time-controlled mode, the algorithm kept track of time and after a preset elapsed time interval the pressurized reservoir was depressurized and *vice-versa*.

Simultaneously, valves changed their position allowing for unidirectional flow. To control fluid flow, a PID controller was employed to counter the sudden flow rate change derived from the valves switching positions. Flow was corrected in the time span of seconds, promptly recovering the preset constant flow (Figure S4). The performance in flow correction of the system here is in the same range as what is found in typical integrated, commercial pumping systems.

Commercial integrated organ-on-chip culture platforms are also capable of recirculating medium, e.g. by integrated peristaltic pumps, or by rocking platforms.^37^ However, such fully integrated ‘closed’ systems are difficult to interface with additional components from other sources. This is understandable for commercial settings where market share is a concern; but for the field as whole, this approach poses limitations on the reusability of the available microfluidic components in the market. The inability to integrate components from different vendors restricts the user to a single company’s product catalogue and forces the re-development of existing devices, delaying application and increasing the entry barrier for new users. By adopting standard interconnections and the FCB concept, users would benefit from a wider range of options while companies would still be able to offer their unique solutions to a bigger market.

The FCB-centered solution for flow control presented here standardizes fluidic connections between various components, thereby also reducing labor while increasing reproducibility. Overall, the system matches the fluidic capabilities of existing commercial solutions while reducing the amount of tubing and connections required.

### Heart-on-chip Culture

To validate the system, we used a HoC device in which three-dimensional cardiac tissues containing human pluripotent stem cell-derived cardiomyocytes were co-cultured with endothelial monolayers of HUVECs, as previously reported ^38^. We were able to use the modular flow control system to fully automate the HoC culture up to 5 days, at which point medium had to be changed manually. HUVECs were able to form a confluent monolayer lining the vascular compartment of the device (Figure 4.a). Three-dimensional cardiac tissues were contracting continuously at an approximate rate of 6 bpm (Figure 4.b, c) and presented a similar morphology to those cultured in control conditions on a rocking platform.

**Figure 4.**
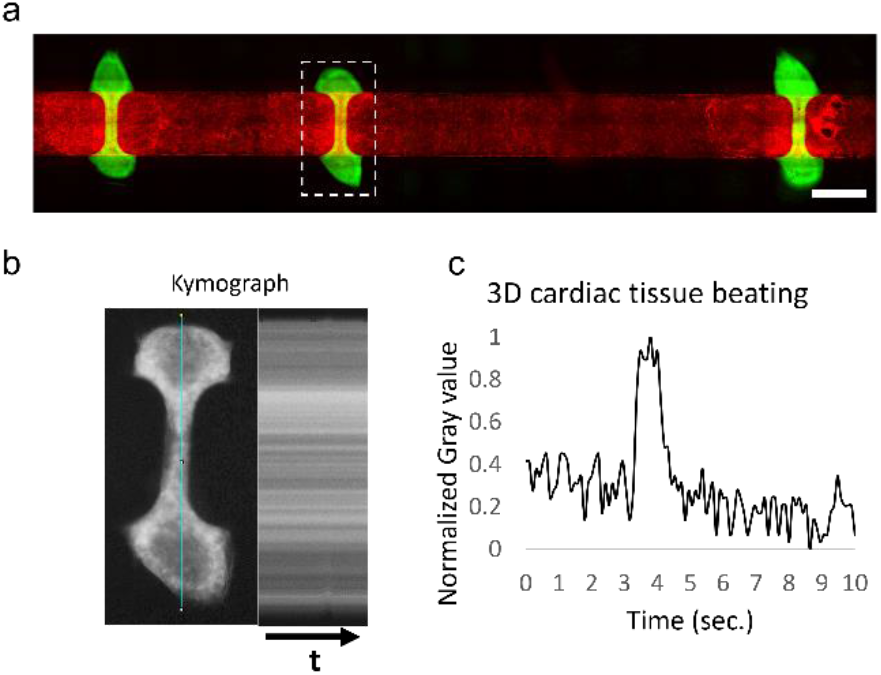
Cell culture in the Heart-on-Chip device, a) Immunofluorescent microscopy images of HUVECs cytoskeleton stained with CellMask™ Deep Red Actin Tracking Stain and 3D cardiac tissues in green by the expression of GFP from the DRAGGN-line. Scale bar: 500 μm. b) Beating frequency graph obtained from a 10 second video clip, right, using Kymography. Left, depicts the tracked line over the duration of the video and right the resulting gray value map over the 10 second period (arrow) of the video, c) Obtained graph from the kymograph detecting the beat of the 3D cardiac tissue.

Despite the continuous shear rate of 833 s^-1^ generated by the medium perfusion, HUVECs did not show any alignment parallel to flow. The conditions on which these cells are cultured are subject to further optimization since the different culture media employed may be altering HUVECs response to shear stress.

## Conclusions

We demonstrated that by using an FCB as a platform to interconnect both commercial and custom-made devices, it is possible to integrate them into a stand-alone solution for organ-on-chip fluidic perfusion. The reduced footprint and tubing of the entire system is enabled by commercially available software and hardware and greatly simplifies the establishment of organ-on-chip culture. We were able to maintain fully perfused organ-on-chip devices with human pluripotent stem cell-derived tissues in culture for up to 5 days in a fully automated fashion.

The presented system is an implementation of the concept of TOP, an open technology platform for organ-on-chip ^16^. The current system highlights the strengths TOP has in facilitating the integration of multiple components into an organ-on-chip culture system that is fit-for-purpose.

Integration of system functionality was achieved by reusing already developed technology both in academic and industrial settings. Systems like the one here presented would allow simplification of fluidic setups in organ-on-chip devices and adoption of a modular approach. Being able to integrate available technologies has the potential to increase adoption of organs-on-chips, since the entry barrier for implementation and customization can be more rapidly lowered. By using an FCB, end-users and developers alike would be able to combine already developed technologies that suit their needs and focus their resources and efforts on the development of their own specific device. Moreover, comparison of results and performance can be more easily attained due to the standardization of connectors and fluidic circuits employed to drive flow through organs-on-chips. Using a standardized approach also enables easier translation from academia to industry settings, since using the same interconnection guidelines would facilitate the coupling of experimental systems to already characterized ones.

We expect that further development and adoption of TOP may help in increased parallelization, compatibility with commercial systems, reusability and collaborations to further develop new organ-on-chip applications.

## Acknowledgements

A.V, A.v.d.B, R.P and A.D.v.d.M acknowledge the European Research Council (ERC) under the Advanced Grant ‘VESCEL’ Program (Grant no.669768) of Prof. Van den Berg.

