## Supplementary Figures for "Fluidic Circuit Board with Modular Sensor and Valves Enables Stand-Alone, Tubeless Microfluidic Flow Control in Organs-on-Chips"

#### Supplementary Figure 1

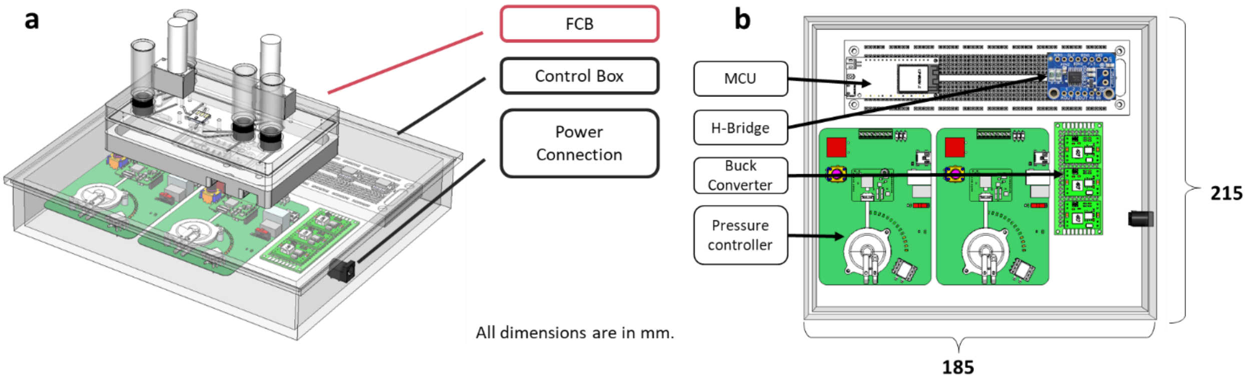


*Supplementary Figure 1. 3D CAD-model of FCB and the control box. A) The FCB and the control box are depicted on top and bottom, respectively. B) the different components are depicted inside the control box, where the layout of the 2 pressure controllers, the MCU, the H-bridge and the buck converters can be seen.*

#### Supplementary Figure 2

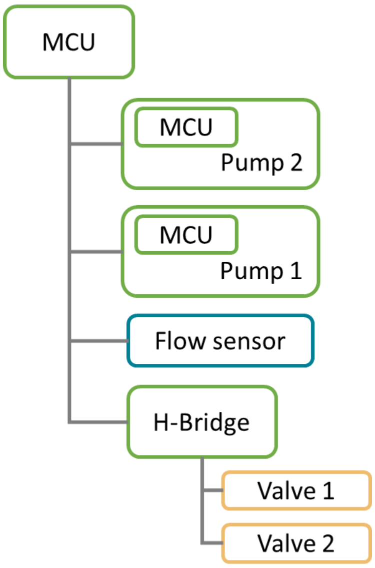


*Supplementary Figure 2. Schematic representation of the electrical connections among the different components. The microcontroller (MCU) is connected with the two pressure controllers (pump 1 and 2). These pressure controllers are composed of a board that contains a microcontroller property of TTP Ventus, a piezoelectric pump. The flow sensor is connected via a ribbon cable and interfaced with the MCU via $I^2C$ protocol. The H-bridge is digitally controlled with two IO pins of the MCU, activating each of the corresponding solenoid valves. These valves are connected to the H-Bridge via a ribbon cable.*

### Supplementary Figure 3

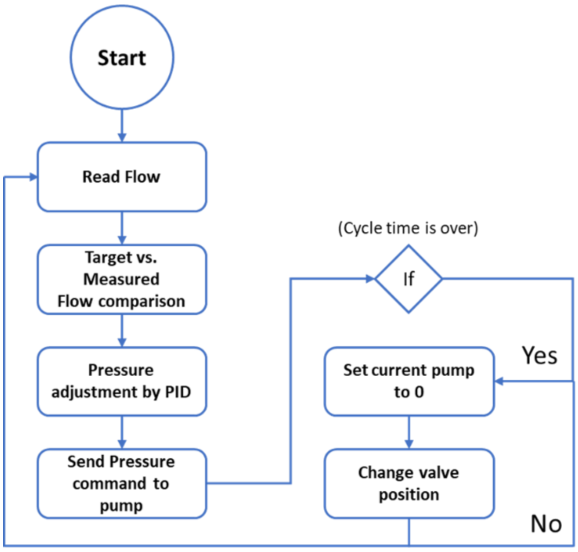


*Supplementary Figure 3. Schematic overview of the control algorithm for flow control. The cycle runs indefinitely until stop by the user and is able to autonomously control the flow rate in the microfluidic device.*

### Supplementary Figure 4

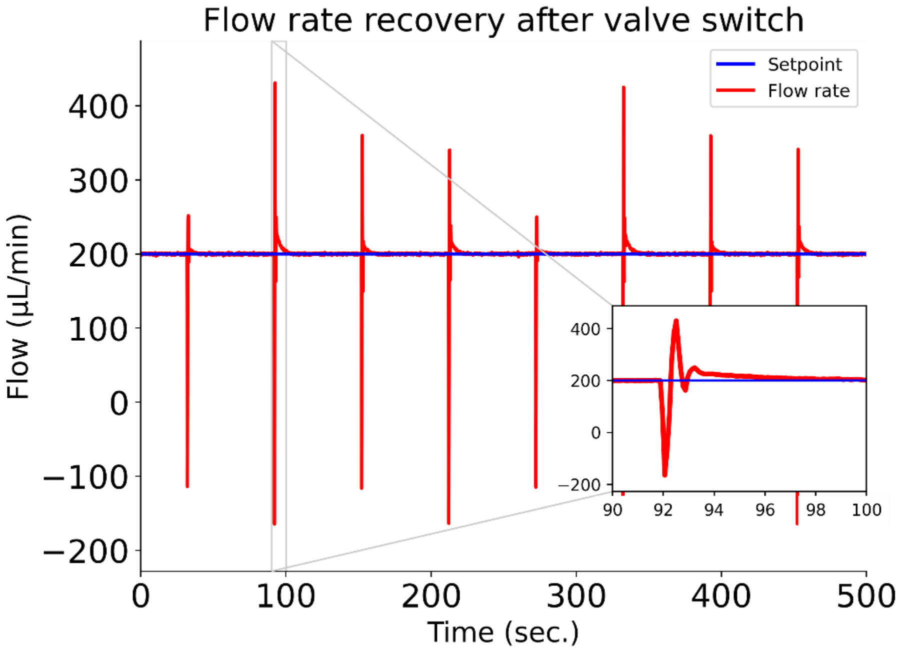


*Supplementary Figure 4. Graph depicting the flow rate recovery after a flow path switch. The system is able to re-establish the set flow rate within seconds of the valves changing positions and reversing flow between the two reservoirs while keeping unidirectional flow in the heart-on-chip vascular compartments.*
